# Cognitive and affective Theory of Mind double dissociation after parietal and temporal lobe tumors

**DOI:** 10.1101/2021.10.26.465856

**Authors:** Fabio Campanella, Thomas West, Corrado Corradi Dell’Acqua, Miran Skrap

**Affiliations:** Neurosurgery Unit, Presidio Ospedaliero Universitario “S. Maria della Misericordia”, Udine, Italy; Department of Life Sciences, University of Trieste, 34100, Trieste, Italy; Faculty of Psychology and Educational Sciences (FPSE), University of Geneva, Geneva, Switzerland

**Keywords:** Brain tumors, Mentalization, Affective Theory of Mind, Cognitive Theory of Mind, Parcel-based Lesion-Symptom Mapping

## Abstract

Extensive neuroimaging literature suggests that understanding others’ thoughts and emotions engages a wide network encompassing parietal, temporal and medial frontal brain areas. However, the causal role played by these regions in social inferential abilities is still unclear. Moreover very little is known about ToM deficits in brain tumours and whether potential anatomical substrates are comparable to those identified in fMRI literature. This study evaluated the performance of 105 tumour patients, before and immediately after brain surgery, on a cartoon-based non-verbal task evaluating Cognitive (Intention Attribution) and Affective (Emotion Attribution) ToM, as well as a non-social control condition (Causal Inference). Across multiple analyses, we found converging evidence of a double dissociation between patients with right superior parietal damage, selectively impaired in Intention Attribution, and those with right antero-medial temporal lesion, exhibiting deficits only in Emotion attribution. Instead, patients with damage to the frontal cortex were impaired in all kinds of inferential processes, including those from the non-social control conditions. Overall, our data provides novel reliable causal evidence of segregation between different aspects of the ToM network from both the cognitive and also the anatomical point of view.

## Introduction

Theory of Mind (ToM) is the ability to attribute and infer one’s own and others’ mental states such as beliefs, expectations and emotions and thereby interpret others’ behavior^1^. Different studies suggest that ToM impairments could lead patients to make faulty judgments regarding other’s cognitive and emotional beliefs^2, 3^, and that this difficulty could lead to social dysfunction but also lower self-perceived quality of life^4–6^. As such, a precise mapping of the neural structures which, if damaged, could lead to ToM deficits is of paramount importance.

ToM is intended as a multidimensional cognitive domain generally including two separate sub-domains: *Affective* ToM, concerning the ability to infer other people’s emotional states, and *Cognitive* ToM, concerning the ability to infer beliefs, intentions and thoughts^7^. From an anatomical point of view, a wide range of studies used brain-imaging techniques to explore the neural foundations of these abilities, and identified partially dissociated mechanisms underlying *Cognitive* and *Affective* aspects of ToM. More specifically, different studies converged in associating *Cognitive* ToM with a widespread network, involving Temporo-parietal junction (TPJ), midde temporal gyrus, precuneus, and lateral and medial PreFrontal cortex (lPFC and mPFC)^8, 9^ (see^10, 11^ for meta-analyses). Importantly, part of this network has been observed also in tasks investigating *Affective* ToM^12, 13^ which, in addition, recruited also the anterior temporal cortex, in both its lateral (temporal pole) and medial (amygdala) aspects, the anterior insula, the inferior frontal gyrus, and parts of mPFC which were not already explained by *Cognitive* ToM processes^14–16^.

Most notably, the precise anatomical localization of the ToM networks has been investigated in a recent meta-analysis on 188 neuroimaging studies^10^, who run Representation Similarity Analysis on meta-analytic brain maps from a wide range of tasks testing different aspects of social cognition. The authors identified three main networks of interest. The first is a *cognitive* network, characterized by TPJ, middle temporal gyrus, precuneus, and mPFC, which was elicited prevalently by *Cogntive* ToM paradigms. The second is an *affective* network, characterized by temporal pole, amygdala, insula, inferior frontal gyrus and middle cingulate cortex. This was elicited by tasks where individuals were exposed with behavioural and somatic consequence of others’ affective states (facial expressions, injured bodies, etc) and, in some cases, were expected to empathize with these states.^17^ The third is a *hybrid* network combining neural structures from the previous two, and elicited by tasks probing abstract reflection and appraisal about others’ emotions. Hence, differently from the purely affective pathway, the hybrid network underlies a more abstract, propositional knowledge how specific contexts and behaviours relate to people emotions/affect^10^, typically employed when inferential mechanisms for ToM are adopted to infer affective states.

In stark contrast with neuroimaging literature, the research investigating the neural underpinnings of ToM from neurological patients is less clear. Indeed, many studies reported deficits in these abilities following neurodegenerative disorders^18–23^ and acquired brain lesions^7, 24–38^. Interestingly, some studies reported dissociations between the neural processes underlying *Cognitive* and *Affective* ToM deficits, for instance in patients with frontotemporal dementia where impairments in the assessment of cognitive and affective states interact differently with syndrome severity or executive functioning^39–42^. Also studies on stroke patients implicated *Cognitive ToM* in the dorsolateral prefrontal cortex^43^, and *Affective* ToM in the ventral mPFC^7^ and limbic regions such as the insula^43^. Unfortunately, the majority of these studies is based on small samples sizes (N ≤ 50), and often include populations with diffuse neural damage^2, 18, 44–46^ or *focal* brain lesions assessed under mixed etiologies (see^47^ as review).

To our knowledge, only few studies investigated the neural underpinning of ToM deficits in large samples. The first^48^ examined ToM abilities in 170 patients affected by penetrating brain lesions and correlated low ToM scores with lesion in the right lPFC and the left parietal cortex, partially confirming evidence coming from fMRI studies. However, this study investigated only *Cognitive* ToM, thus leaving open the question as to whether the implicated regions are necessary for inferring also other kinds of states (e.g., emotions).

Further studies examined brain tumour^28, 49, 50^ and stroke patients^29^ (samples ranging between 64-122 patients) on a ToM task requiring to infer others’ states from gazes (Reading the Mind in the Eyes Test; RMET^51^). These studies converge in highlighting a critical role of TPJ^28^, middle temporal cortex^28^, posterior frontal gyrus^29^, and insula^29, 28^, as well as of the white matter tracts linking frontal and temporo-parietal areas^29 49 50^. In one case, authors found a dissociation between RMET, and similar Comic Strip task (derived from^52^) aimed at testing similar ToM abilities under more abstract, high-level, inferential processes^49^. Importantly, however, in all these studies the absence of two clear-cut *Cognitive* vs. *Affective* conditions did not allow to disentangle appropriately the neural correlates of these ToM components. Furthermore, in some cases, the lesion distribution was substantially biased towards the frontal lobe thus providing insufficient coverage of all relevant brains structures^49 50^.

From a more clinical point of view, in spite of a large number of experimental tasks developed evaluating single processes involved in ToM, only a few *clinically validated* tools are available to assess the integrity of both cognitive and affective domains^6^. Moreover, the most frequently used tasks in clinical settings involve verbal materials (e.g., text-based storyboards) which are vulnerable to linguistic confounds. Other paradigms are challenging due to their length or complexity (e.g. with high loadings on executive functions or working memory), reducing the feasibility of ToM assessment in most severe clinical populations, and limiting the generalizability of results regarding the integrity of “core” ToM components.

The present work aims at overcoming the limitations of previous research, by investigating for the first time the neural underpinnings of *Cognitive* and *Affective* ToM deficits in a large population of 105 brain tumor patients, through a validated clinical task (the Story-Based Empathy task: SET^53^) which does not rely on verbal materials and with minimal working memory demands. Although originally classified as an “empathy” task, the SET requires individuals to infer from a comic-strip a character’s intentions (Intention Attribution [IA] condition) or emotions (Emotion Attribution [EA]), thus probing individual ToM abilities applied to both cognitive and affective states.

Based on the literature reviewed above, we expected to find stable lesion correlates to these abilities around the parietal^28, 49^, temporal, limbic^28^ and prefrontal cortex^48, 49^. The critical question, however, is to which degree these dissociate from one another and, in case, whether this dissociation is consistent with that observed in neuroimaging literature on neurotypical individuals^10^, thus allowing to identify components of the ToM network which are necessary for different kinds of social inference.

## Materials And Methods

### Participants

105 patients undergoing surgery for the ablation of brain tumors participated to the study. Inclusion criteria were: being affected by gliomas or meningiomas, having a sufficient clinical and/or cognitive picture (i.e. being able to undergo a complete neuropsychological evaluation), being aged between 18 and 75, undergoing first surgery. Patients with multifocal as well as recurring lesions were excluded. Participants underwent a complete neuropsychological evaluation covering main cognitive domains (language, attention, executive functioning, memory and perception) a few days before surgery and 5-7 days after. Table 1 provides full details demographic characteristics of patients’ sample organized based on the lobe location. Age and Education distribution was comparable across hemispheres and lobe locations (See Table 1), while Lesion Volume was found to be higher for right hemisphere patients. The study was approved by the ethical committee of A.S.U.I. Udine and patients provided written consent.

**Table 1.** Demographic and histological characteristics of the sample of patients included in the study divided according to lesion location. Bold indicates significant differences ^a^ = χ^2^ test; ^b^ = Independent samples T-test; ^c^ = One-Way ANOVA.

|  | Lateralization |  |  | Location |  |  |  |
| --- | --- | --- | --- | --- | --- | --- | --- |
|  | Left | Right |  | Frontal | Temporal | Parieto-Occipital |  |
| Number of patients (n=105) | 55 | 50 | p=.782<sup>a</sup> | 49 | 34 | 22 | p=.070<sup>a</sup> |
| Mean Age: |  |  |  |  |  |  |  |
| Years (SD) | 53.18 (14.82) | 55.02 (14.75) | p=.394<sup>b</sup> | 55.14 (15.73) | 54.27 (12.72) | 51.32 (15.70) | p=.601<sup>c</sup> |
| Mean Education: |  |  |  |  |  |  |  |
| Years (SD) | 12.15 (3.86) | 12.44 (3.64) | p=.526<sup>b</sup> | 11.78 (3.75) | 12.71 (3.71) | 12.77 (3.82) | p=.430<sup>c</sup> |
| Mean Lesion Volume: |  |  |  |  |  |  |  |
| Cm<sup>3</sup> (SD) | **76.42 (65.53)** | **112.35 (83.90)** | **p=.014<sup>b</sup>** | 85.28 (85.72) | 102.95 (63.37) | 97.32 (68.60) | p=.560<sup>c</sup> |
| Tumor Hystology |  |  |  |  |  |  |  |
| HGG (n=43) | 20 | 23 |  | 12 | 19 | 12 |  |
| LGG (n=32) | 12 | 11 |  | 11 | 8 | 4 |  |
| Meningioma (n=39) | 23 | 16 |  | 26 | 7 | 6 |  |
|  | p=.386<sup>a</sup> | p=.332<sup>a</sup> |  | p=.144<sup>a</sup> | p=.175<sup>a</sup> | p=.335<sup>a</sup> |  |

### Theory of Mind assessment

We evaluated of Cognitive and Affective Theory of Mind skills by means of the Story-based Empathy Task (SET)^53^, administered before and also after surgery. The SET is a completely non-verbal task in which patients are required to select the correct ending of a comic strip, among three possible endings provided below the main story (see Fig.1). The task consists of two Theory of Mind conditions, i.e., Intention Attribution (IA) and Emotion Attribution (EA), where participants inference is based upon the correct interpretation of the intention or the emotional state of the main character respectively. In a further control condition (Causal inference, CI) the correct answer is based on the assessment of physical properties of the world, which controls for more general impairments in basic reasoning skills^53^. Each condition includes six trials (see Supplementary Material for further details). No verbal answer or verbal processing is required at any stage and no facial expression is depicted in any of the vignettes which may influence the assessment; all the materials necessary for comic strips interpretation and answer is constantly in sight of the patient thus drastically reducing the working memory load.

**Fig. 1:**
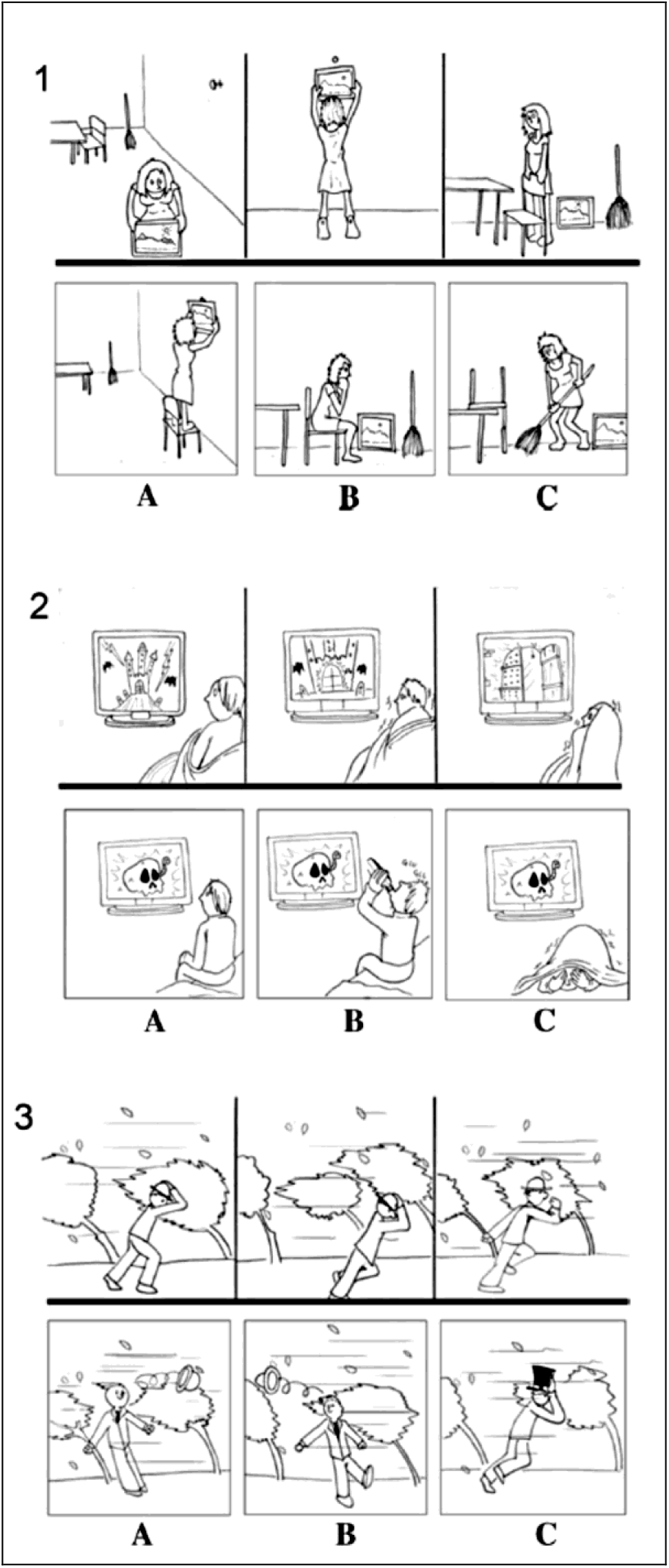
Vignettes from the Story-based Empathy Task. 1) Intention attribution (SET-IA), 2) Emotion attribution (SET-EA) based on fear, 3) causal inference (control condition—SET-CI). A, B and C represent the possible endings of the story among which subjects must choose the correct one. (Reproduced from Dodich et al, 2015).

### Behavioral data analysis

All raw scores in each condition were corrected for age and education and transformed into z-scores based on standardization data^53^. As first step, we subdivided the population into three anatomical groups: Frontal, Temporal and Parieto/occipital. Patients were assigned to each lobe location according to the predominant lobe of involvement of the lesion. This was assessed by superimposing the normalized MRI scan to the standard AAL (Automated Anatommical Labeling^54^) template and calculating the precise percentage of lesion involvement for each sub- region of each lobe to compute which lobe was more involved.

#### ANCOVA Analysis

An ANCOVA was performed on the whole patient group considering “Hemisphere” (Left vs. Right), “Lobe Location” (Frontal vs. Temporal vs. Parieto/occipital) as between-subject variables and “Surgery” (Pre vs. Post-surgery) and “ToM Condition” (CI vs IA vs. EA) as within-subject factors, to evaluate whether specific anatomical regions were linked to lower scores in any of the experimental conditions. Age, education and lesion volume were added as covariates whereas “Etiology” (HGG vs. LGG vs. Meningioma) was added as a between-subject nuisance factor. Within this analysis, we modeled all main effects, and all interactions between the within-subject factors and grouping factors: Hemisphere, Lobe Location, Hemisphere*Lobe Location, and all nuisance variables. This allowed to assess whether the critical “ToM Condition” factor interacted with the anatomical location of the lesion (described by Lobe Location/Hemisphere) while accounting for non-specific grouping confounds. Post-hoc analysis was then performed to investigate the sources of possible significant main effects and interaction, by means of Tukey Test. Sensitivity analysis over the design revealed that the ANCOVA could reliably detect effects associated with the ToM Condition within-subjects factor ranging around 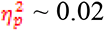, under power = 0.8 and α = 0.05 (see Supplementary Information for more details).

### Clinical evaluation: incidence of ToM deficits in the sample of patients

As follow-up, we assessed whether the average z-score obtained by each anatomical subgroup of patients (Frontal, Temporal and Parieto-Occipital) in each condition of the task was significantly lower than expected from healthy population (based on standardization cut-offs^53^), in order to evaluate the occurrence of ToM *deficits*. The analysis was performed by means of a series of one-sample T-Tests, against an expected average z-score of 0 with a standard deviation of 1. Bonferroni correction was adopted to correct for multiple comparisons between ToM conditions (p=0.05/9=0.006) for each anatomical subgroup.

### Lesion reconstruction and anatomical PLSM analyses

For all patients, pre-operatory high resolution gadolinium-enhanced T1 and (when available) T2-weighted and/or FLAIR MRI scans for neuronavigation (minimum number of slices: 180) were collected to determine tumor location. The 3D region of interest (ROI) reconstructions of lesions were drawn for each patient from MRI slices on the horizontal plane using MRIcroN software^55^. ROIs included all altered MRI signal areas, including edema, which is known to have cognitive effects^56–59^. To match and align images on a common Montreal Neurological Institute (MNI) template, each MRI scan then underwent spatial normalization using SPM12 (https://www.fil.ion.ucl.ac.uk/spm/) software. Lesion volumes were thus calculated from normalized ROIs.

Anatomical lesion-symptom mapping analyses were then performed by means of Parcel-based Lesion-Symptom Mapping (PLSM), to better specify anatomical regions most critically linked to a lower performance in each ToM condition. The brain was divided in 116 regions of interest (ROIs) from the AAL grey-matter atlas^54^, implemented in the NiiStat toolbox for Matlab (http://www.nitrc.org/projects/niistat). This allowed the investigation of gray matter correlates most likely linked to potential ToM deficits. PLSM was performed only in those voxels damaged in at least 6 patients, as the best compromise between having a “sufficient lesion affection”^60^ and the need to sample a sufficiently wide and homogeneous brain surface (see Supplementary Fig.1). Family-wise error rate was controlled via 5000 permutations, with a threshold set at p < .05. A Monte Carlo simulation was run to assess the sensitivity of each ROI to detect effects under power = 0.8 and α = 0.05 corrected for multiple comparisons, and reveal that all the regions included could detect effects of at least *r* ∼ 0.39 (see Supplementary Information for more details). Since no effect of surgery was found in any condition (see Results) for the PLSM analysis pre-post surgery scores were averaged and used as dependent variables. To investigate the anatomical substrates of both Emotion (Affective ToM) and Intention (Cognitive ToM) Attribution, a separate PLSM analysis was performed on each of the two conditions.

#### PLSM analysis 1: areas linked to worse ToM deficits (EA<CI and IA<CI)

Scores considered as dependent variable were the result of the subtraction between the z-scores obtained in the CI condition (control condition) and those obtained in EA and IA conditions separately, in order to highlight only those anatomical areas linked specifically to ToM processes cleared from the influence of any potential more general “causal” or “logical” reasoning difficulty. Furthermore, these differential scores were subsequently modeled as function of all nuisance variables who had a significant effect in the main ANCOVA (i.e. Age, Education and Lesion Volume, see Results). The residuals from this linear model were then fed to the PLSM procedure. This insured to identify lesion-symptom associations independently from non-specific confounds.

#### PLSM analysis 2: areas linked to greater ToM selectivity (IA<EA and EA<IA)

A second PLSM analysis was then performed to highlight potential anatomical regions in which, apart from lower scores in general, the *gap* between the performance in the two ToM conditions (IA and EA) was larger (i.e. the selectivity of the effect) using as dependent variable the differential score between IA *vs*. EA directly. Also in this case, potential confounds of Age, Education and Lesion Volume were removed from these differential scores.

### Data availability statement

The data that support the findings of this study are available from the corresponding author upon reasonable request.

## RESULTS

### Behavioural results

#### ANCOVA analysis

Group analysis revealed no main effect of Hemisphere, Lobe Location or ToM Condition over patients’ performance on the task (see Supplementary Table 1 for full results). Most interestingly, however, we found a highly significant interaction between ToM Condition and Lobe Location (F_(4,168)_=5.907 p<.001; η^2^=.123) suggesting that performance in the three ToM conditions was differentially influenced by the lobe location of the lesion. Post-hoc analysis showed that Parieto/Occipital Lobe lesions (regardless of the lateralization) led to greater difficulties in Intention Attribution (IA), with respect to both Causal Inference (CI; p=.015) and Emotion Attribution (EA; p=.002; see Fig. 2). On the contrary, in clear dissociation with this, Temporal Lobe lesions led to an opposite pattern of performance, with patient showing selectively worse performance on EA with respect to CI (p=.009) and marginally to IA (p=.055). Patients with Frontal Lobes lesions instead did not show any condition difference, with the exception of a marginal discrepancy between IA and CI (p=0.054). Regarding covariate influence, a significant main effect of Age F_(1,84)_=9.987; p=.002; η^2^=.106), Education F_(1,84)_=5.397; p=.023; η^2^=.060) and Lesion Volume (F_(1,84)_=5.747; p=.019; η^2^=.064) was found, indicating that older and less educated patients and with larger lesions performed generally worse on the task. A significant interaction was further found between both Education (F_(2,168)_=4.823; p=.009 η^2^=.054) and the ToM condition, basically showing that less educated patients scored worse in both IA and EA but not CI (i.e. they were more prone to ToM deficits). No significant effect or interactions were found for the nuisance factors Etiology (type of tumor) and Surgery effects (before vs. after ablation), neither when considering main effects nor interactions with other variables (see Supplementary Table 1). Given the lack of surgery effects, all further analyses from here on, were performed on the average score between pre- and post-operatory performance of each patient.

**Fig. 2:**
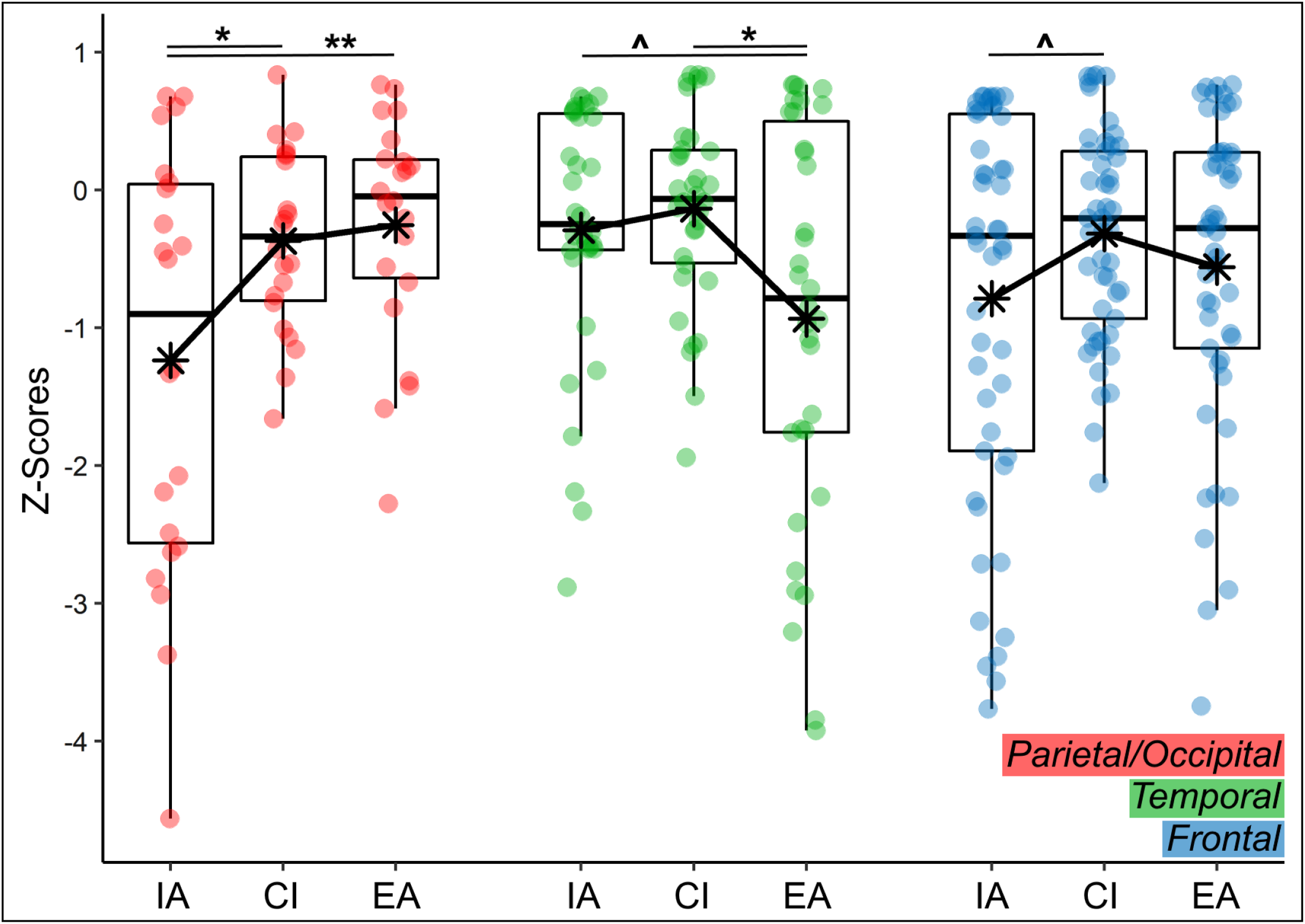
SET task behavioural results: patients with Frontal and Parieto/Occipital lesions had greater difficulties in interpreting intentions, while Temporal patients had greater difficulties in interpreting emotions. IA=Intention Attribution; CI= Causal Inference; EA= Emotion Attribution. **^**=p=.055; *=p<.05; **=p<.01;

#### Clinical evaluation: incidence of ToM deficits in the sample of patients

Table 2 displays full results from the clinical evaluation, confirming that the pattern of performance found in the ANCOVA was reflected also in terms of incidence of ToM *deficits*. In particular, Frontal Lobe patients had significant deficits in both IA and EA, but also in CI. Parieto/Occipital Lobe patients instead had deficits only in IA, while, on the contrary, Temporal Lobe patients had deficits only in EA, both before and after surgery. Thus, clinical results seem to suggest that Frontal Lobe patients had more diffuse ToM deficits, but that these might be influenced by also a more general difficulty in abstract reasoning/executive functioning.

**Table 2.**
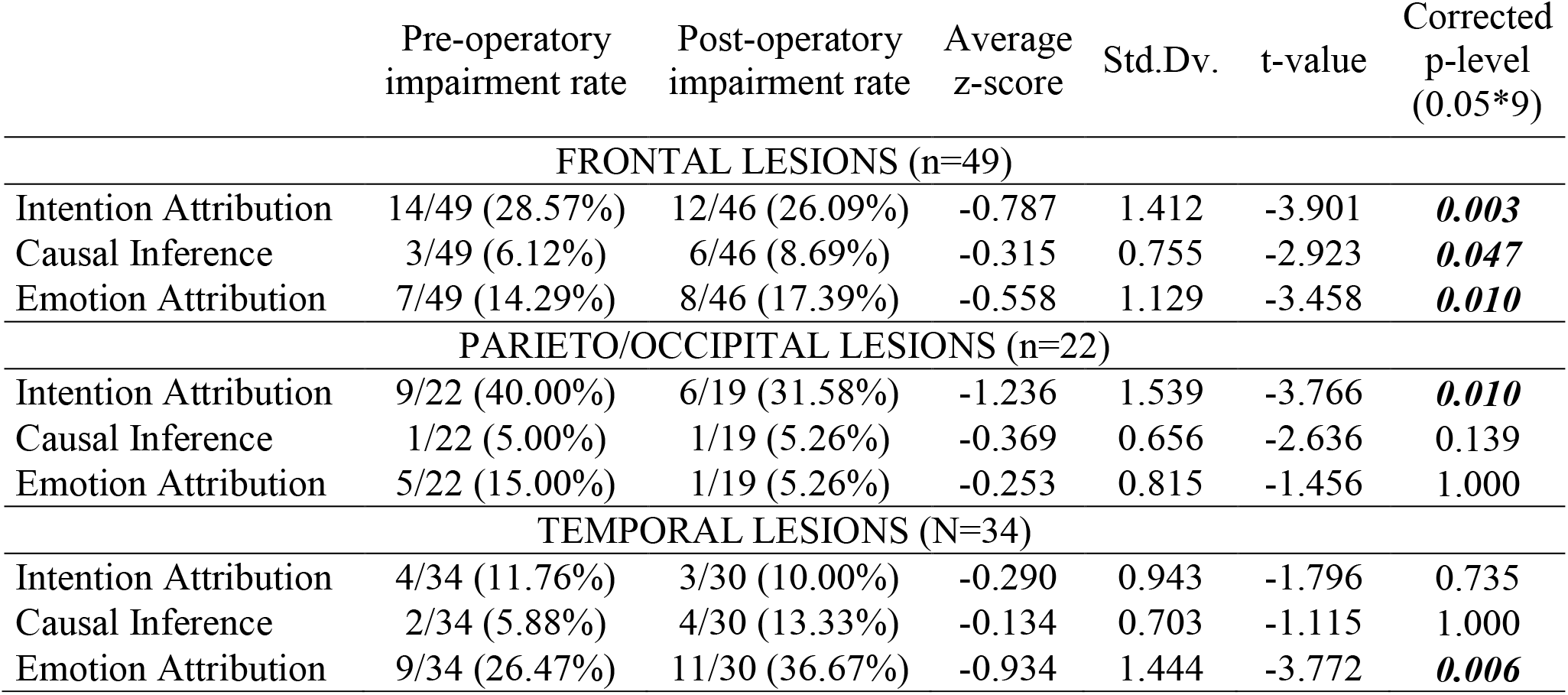
Clinical summary: performance of patients in the Story-based Empathy Task (SET), with number of patients (and %) obtaining clinically relevant (at least borderline) scores in each condition before and after surgery. One-sample T-test results indicate groups performing significantly worse than expected. P-levels were Bonferroni corrected for multiple comparisons within each condition (p-level x 9). Bold indicates significant contrast after correction.

|  | Pre-operative<br>impairment rate | Post-operative<br>impairment rate | Average<br>z-score | Std.Dv. | t-value | Corrected<br>p-level<br>(0.05*9) |
| --- | --- | --- | --- | --- | --- | --- |
| FRONTAL LESIONS (n=49) |  |  |  |  |  |  |
| Intention Attribution | 14/49 (28.57%) | 12/46 (26.09%) | -0.787 | 1.412 | -3.901 | **0.003** |
| Causal Inference | 3/49 (6.12%) | 6/46 (8.69%) | -0.315 | 0.755 | -2.923 | **0.047** |
| Emotion Attribution | 7/49 (14.29%) | 8/46 (17.39%) | -0.558 | 1.129 | -3.458 | **0.010** |
| PARIETO/OCCIPITAL LESIONS (n=22) |  |  |  |  |  |  |
| Intention Attribution | 9/22 (40.00%) | 6/19 (31.58%) | -1.236 | 1.539 | -3.766 | **0.010** |
| Causal Inference | 1/22 (5.00%) | 1/19 (5.26%) | -0.369 | 0.656 | -2.636 | 0.139 |
| Emotion Attribution | 5/22 (15.00%) | 1/19 (5.26%) | -0.253 | 0.815 | -1.456 | 1.000 |
| TEMPORAL LESIONS (N=34) |  |  |  |  |  |  |
| Intention Attribution | 4/34 (11.76%) | 3/30 (10.00%) | -0.290 | 0.943 | -1.796 | 0.735 |
| Causal Inference | 2/34 (5.88%) | 4/30 (13.33%) | -0.134 | 0.703 | -1.115 | 1.000 |
| Emotion Attribution | 9/34 (26.47%) | 11/30 (36.67%) | -0.934 | 1.444 | -3.772 | **0.006** |

To further investigate this possible link, the performance of patients in each of the ToM conditions (IA and EA) was correlated with their attention/executive functions level. This score was obtained by averaging the z-scores obtained by each patient in the Trail Making Test (condition B-A), a measure of attentional switching^61^ and in the Weigl Sorting Test^62^, a test of executive functions measuring abstract reasoning and categorization abilities (a kind of “simplified” version of the Wisconsin Card Sorting Test), which were included in routine neuropsychological evaluation of all patients. As is evident from the scatterplot in Fig.3, there was a strong relationship between executive function level and IA and EA performance for patients with Frontal Lobe lesion, both when taking into account scores from before and after the surgery. This was not the case for patients with either Temporal or Parietal Lobe lesions for which no positive relationship was identified.

**Fig. 3:**
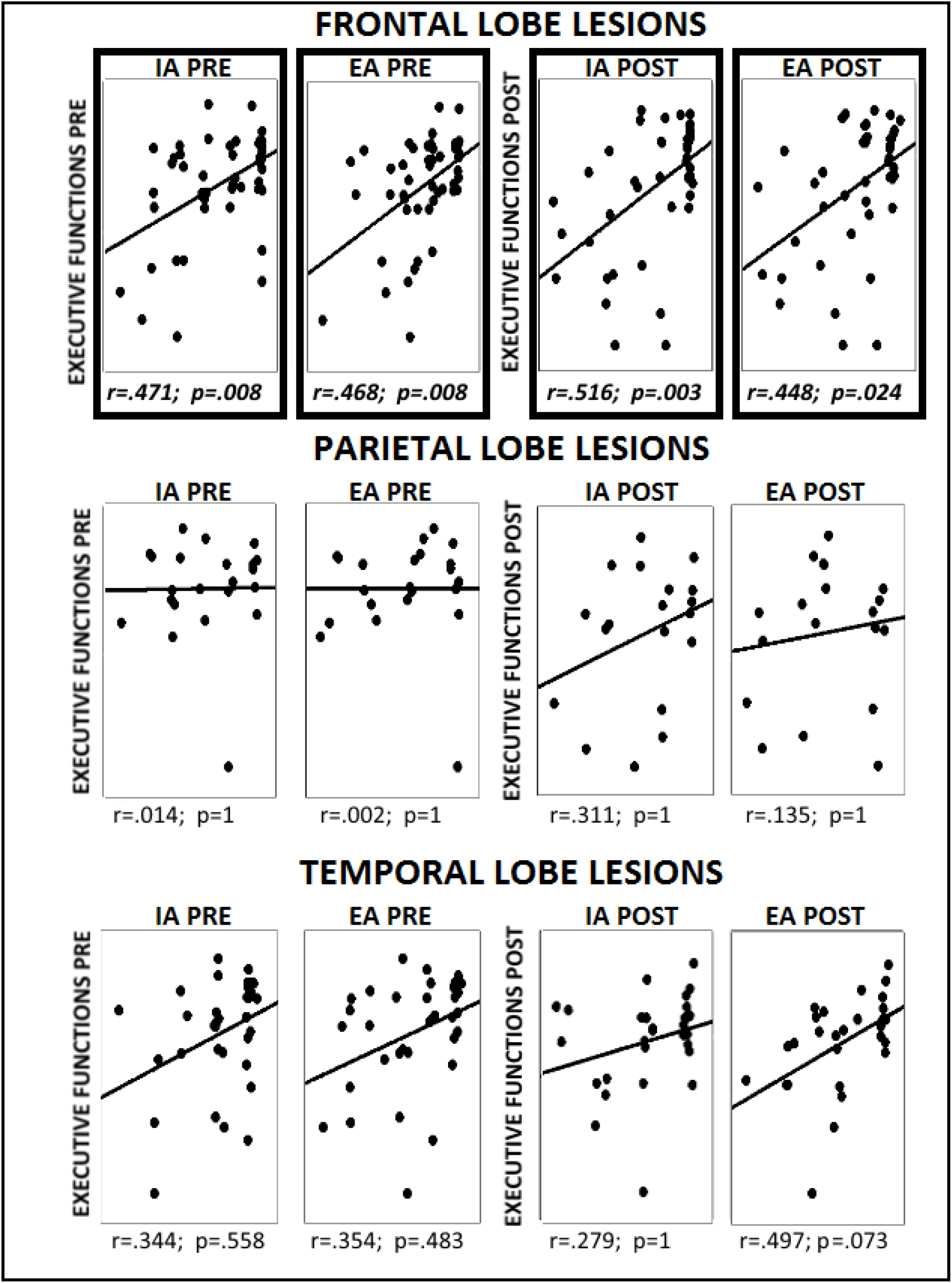
performance of Frontal Lobe patients in Cognitive and Affective ToM conditions was associated with their executive functions level, while for Parietal and Temporal patients, ToM and abstract reasoning levels appeared to be independent. P-level was Bonferroni-corrected for multiple comparisons (p= .05/12 =.004)

A final control was made, with the same rationale, to check whether basic mood state was in some way correlated with IA or EA scores in the three groups of patients (see Supplementary table 3). No correlation was found either before or after surgery with IA or EA scores for both state anxiety (STAI-Y1) or depressive state (BDI-II) measures (all p>0.105).

### Anatomical PLSM Results

#### PLSM analysis 1: areas linked to worse ToM deficits (EA<CI and IA<CI)

Results of the first PLSM analysis performed over the IA and EA scores are shown in Fig.4. A significant cluster of association was found in the right Amygdala (z=-3.91) and Right Superior (z=-3.77) and Middle (z=-3.36) Temporal Pole, for EA scores (relative to CI). No significant association with frontal lobe lesions was found for EA. Furthermore, no significant anatomical correlate was also found for lower IA scores in general, either in the parietal or frontal areas (or elsewhere).

**Fig. 4:**
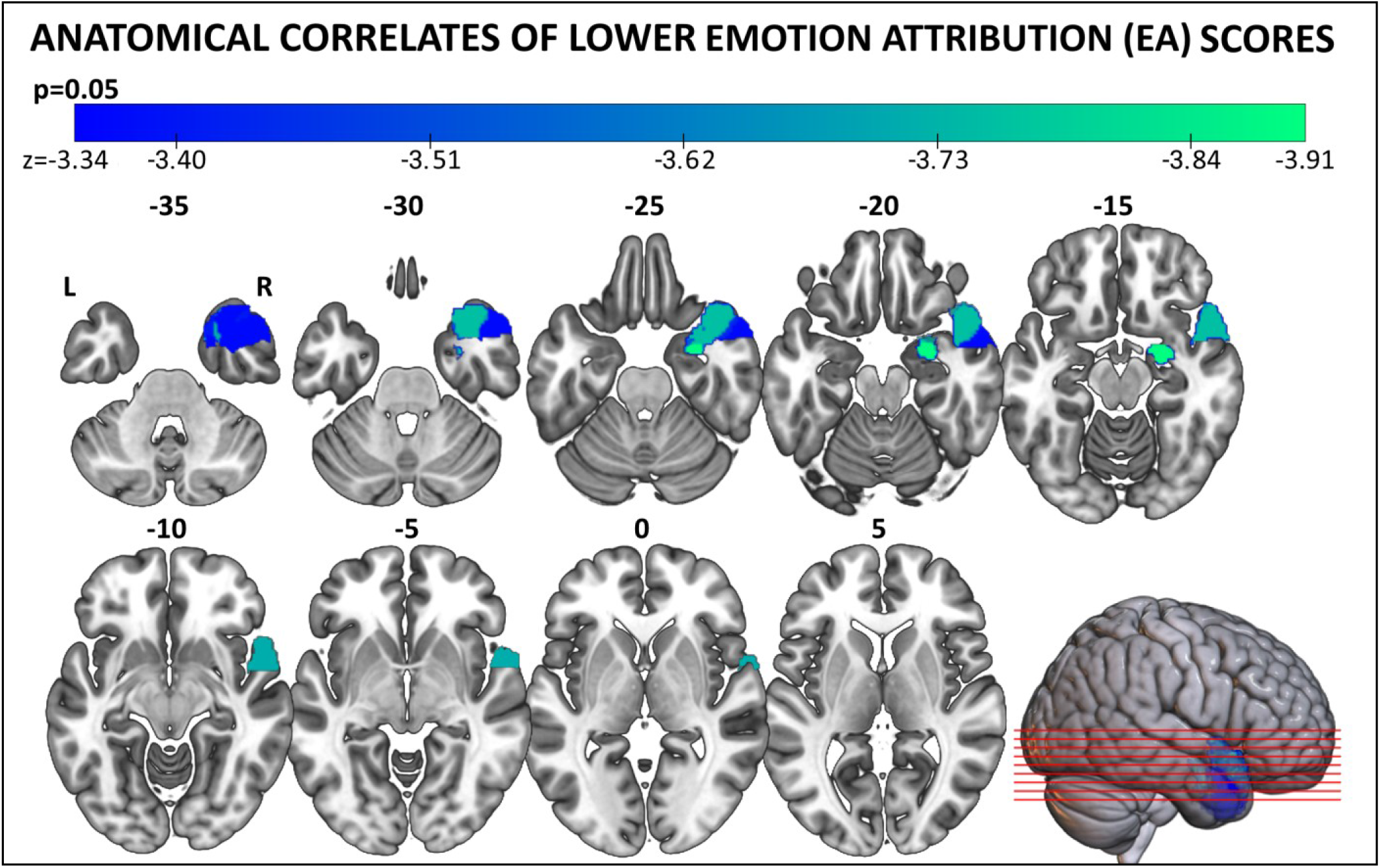
Right Anterior Temporal Lobe lesions (Amygdala and Superior and Middle Temporal Pole) were associated with lower Emotion Attribution (EA) scores. No specific region was associated with generally lower scores in Intention Attribution (IA).

#### PLSM analysis 2: areas linked to greater ToM selectivity (IA<EA and EA<IA)

The results of the second PLSM analysis, for the selectivity of the effects, are shown in Fig.5: in the Right Amygdala (z= -3.450) the gap between EA and IA scores was larger, with worse EA scores, confirming the effects from the previous analysis. Conversely, another significant cluster of association was found in the Right Superior Parietal Lobe (r-SPL; z=-3.251), in which the gap between IA and EA scores was larger, but with worse IA scores. Again no involvement of frontal lobe regions was found also at this level of analysis for any of the scores considered.

**Fig. 5:**
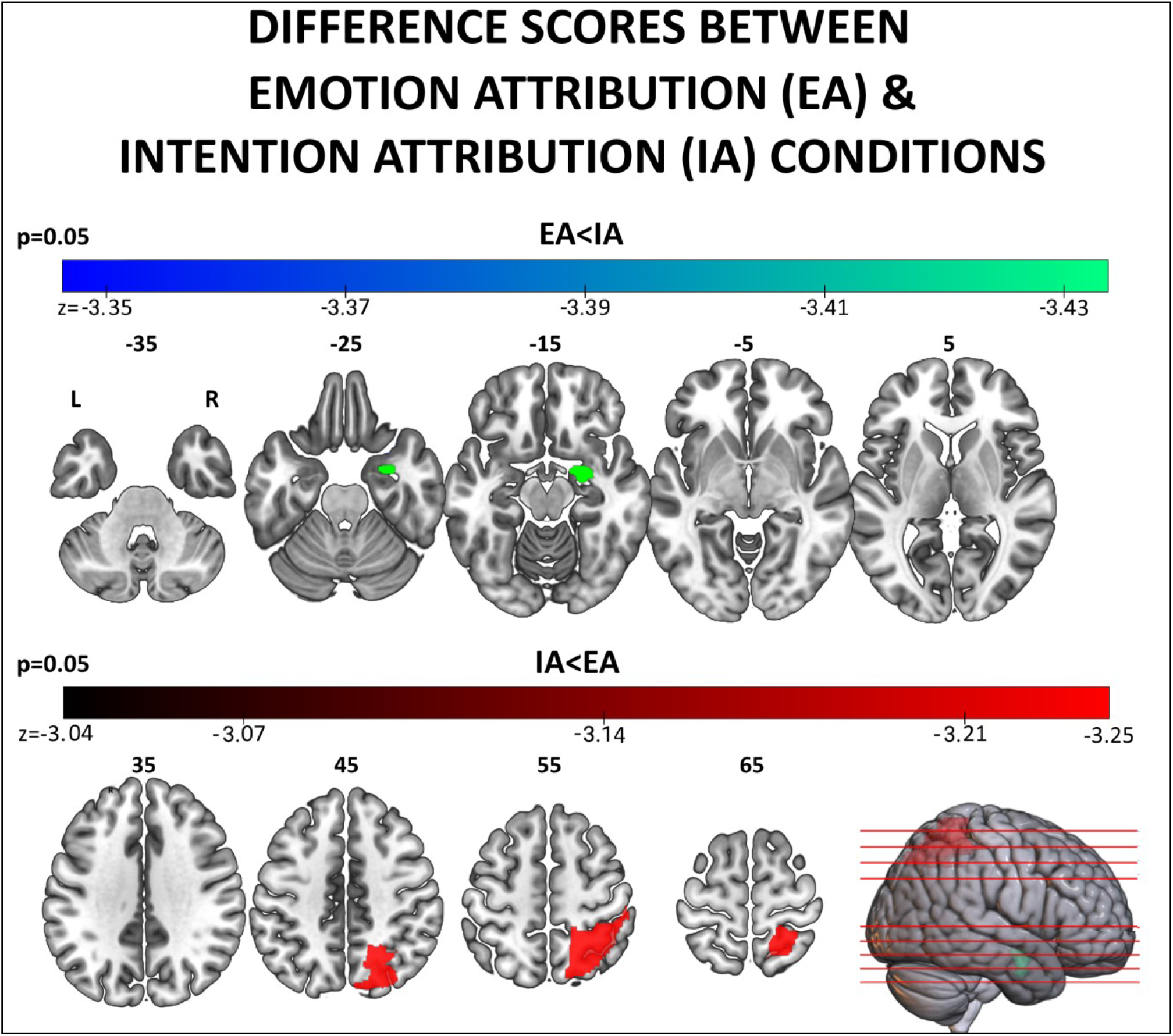
Difference scores between EA and IA conditions: in the Right Superior Parietal lobe IA scores were particularly worse than EA scores. The opposite dissociation was found in the Right Amygdala

## Discussion

We found a striking dissociation in the information represented within the human brain structures mediating inferential abilities about others’ cognitive and affective states. By evaluating a large population of brain-damaged patients (n=105) with an unbiased task for clinical ToM assessment^53^, we provide novel evidence that *Cognitive* ToM deficits are selectively associated with damage to the right parietal cortex, whereas impairments in inferring others’ affect are linked to damage right temporal pole and amygdala. Finally, deficits in both aspects of ToM are associated with damage to the frontal cortex, although such effect could be influenced by broad inferential abilities that generalize also to non-social contexts, and is modulated by patients’ executive impairments. Overall, our study overcomes the limitations of previous research based on populations with low power (see^21^ for review), or investigating only one ToM aspect^28, 48–50^ ^29^ by reliably estimating the anatomical correlates of *Cognitive* and *Affective* metalizing abilities, and by revealing that these are only partly similar to those observed in neuroimaging research on neurotypical individuals.

### Clinical evaluation and behavioural results

Our study underlines how ToM deficits in Brain tumour patients are frequent, and that the evaluation of these processes should be included in routine neuropsychological assessments (see also^6^). In this perspective, the SET task proved to be a very effective clinical tool, due to its feasibility, and limited load on patients’ linguistic, memory and attentional abilities. The only instance in which SET performance appeared to be influenced by processes unrelated to social inferential abilities, was in the analysis of frontal lobe patients. In general accordance with previous literature^7, 43, 48^, frontal lesions led to significant ToM deficits (see Table 2). However, these same patients experienced significant problems also in the Causal Inference control condition, suggesting that their deficit might be influenced/exacerbated by a more general difficulty in abstract-causal reasoning^63^. This was further corroborated by a subsequent correlation analysis revealing how, only in Frontal Lobe patients, there was a positive relationship between task performance and executive function/abstract reasoning skills.

In striking contrast with the frontal cortex, damage to temporal and parietal lobes led to much more selective ToM deficits. Indeed, patients with temporal lobe lesions showed deficits *prevalently* in the *Affective* ToM score, whereas patients with parietal lobe lesions were impaired *prevalently* in the *Cognitive* ToM condition. Within each group there were few cases with deficits in both conditions (see Fig. 2), but even in these cases the deficits were more pronounced for *Affective ToM* in temporal patients, and *Cognitive ToM* in parietal cases. This represents a clear double dissociation^64, 65^, which is immune to confounds from either the population (e.g., age, education, lesion size, tumor kind) or the task (IA, EA and CI shared similar characteristics^53^).

Critically, our results could not be influenced by patients’ linguistic proficiency, since the SET is completely non-verbal and can therefore be easily performed also by aphasic patients. Similarly (and differently from many other ToM tasks), SET task has virtually no load on Working Memory, as all of the elements necessary to infer the correct answer are available to the patient at the same time. Finally, in the *Affective* ToM condition, no influence of more basic emotion recognition skills can be considered as influencing the results, as facial expressions are never displayed, and people’s affective state can be accessed only through the correct interpretation of the situation.

An unexpected outcome from our study was the absence of any surgery effect in the SET. Brain surgery usually leads to additional deficits as consequence of the ablation of functioning brain tissue surrounding the lesion. This is more likely in LGG (in which, when possible, a supra-total resection is always attempted), but less likely in HGG (in which only the compact and often necrotic part of the lesions is usually ablated) and Meningiomas (extra-axial lesions) which constitute the large part of the present sample of patients. However, detrimental surgery consequences might be counterbalanced by the opposite beneficial effects of pressure relief over the immediate neighboring tissue. Please notice, however, that surgery had an impact on other scores from the neuropsychological assessment, with patients exhibiting a decline in linguistic proficiency following ablation of part of the temporal lobe (see Supplementary Table 2 for full details). It is unclear why SET performance was immune to surgery, which appears in contradiction with previous research testing ToM abilities though the RMET^66, 67^. Interestingly however, the RMET is a highly idiosyncratic ToM paradigm, as it relies on direct exposure of facial parts, thus sharing more similarities with many emotion recognition tasks (see^10^ for a similar argument). Indeed, studies finding surgery effects on the RMET, report also no decline in other paradigms requiring abstract appraisal of cognitive/affective states in absence of overt display of facial or body movements: these include Alexithymia tests^66^ Comic Strip Task for the assessment of ToM which bares many similarities with the one used here^67^. In this perspective, it is possible that high-level abstract inferential abilities, which are less “encapsulated” in one specific system and rely in larger extend on distributed network activity, might be less vulnerable to surgery effects. Alternatively, it is in principle possible that surgery effects might have been mitigated by practice, as the same task was repeated twice in short time. Unfortunately, we cannot exclude this possibility since the SET does not have an alternative version and we did not have a control group for testing for test-retest effects. It should be mentioned, however, that other tasks from the neuropsychological assessment, who were repeated in identical form within the same short period, exhibited declined scores following surgery (see Supplementary Table 3). Hence, at least for these cases, practice effects were not sufficiently strong to mitigate surgery-related decline.

### Parcel-based Lesion-Symptom Mapping

PLSM results indicated two clusters of areas (anterior temporal pole/amygdala and superior parietal lobe) significantly linked to difficulties in both *affective* and *cognitive* ToM. Both these clusters were right lateralized. These results are in good agreement with previous evidence coming from lesion studies, frequently reporting right temporal/insular damage in many social cognition tasks, testing empathy^21, 68^, affective ToM^31^ but also investigating more broad ToM abilities^6, 28, 69^

#### Emotion Attribution (Affective ToM)

Anatomical results indicated that damage to the lateral (temporal pole) and medial (amygdala) portions of the right anterior temporal cortex are selectively associated with impairments at reasoning about others’ emotional states. These results are in keeping with previous studies using the SET on patients suffering from the behavioural variant of Fronto-Temporal Dementia^70, 71^, in which significant deficits in the Emotion Attribution condition were consistently correlated with damage to the right Amygdala and right temporal pole (but see^72^ for deficits also in cognitive ToM following the same local atrophy). Furthermore, these same regions have also been extensively implicated in neuroimaging studies in neurotypical individuals, at the point that the PLSM clusters found in our study overlap nicely with previous meta-analytic work from Schurz and colleagues^10^, specifically with the ‘hybrid’ map for cognitive/affective social processing, especially in relation to the ‘reasoning about emotions’ task (see Fig. 6). One reasonable interpretation is that this network stands at the crossroads between the mechanisms involved in pure affective processing and those implicated in social inferential abilities, sharing aspects with both but also displaying unique properties. In this view, some regions might display common neural mechanism between *cognitive* and *affective* ToM, underlying the exploitation of a unique core inferential process (e.g., this was as suggested for TPJ^16, 43^ see below). Other regions, instead, might be tasked to bridge the output of such inferential process with a representation of selective affective states, and therefore might be uniquely implicated in *affective ToM*. Based on our results, the temporal pole and amygdala are two obvious candidate regions for the latter process. Indeed, not only they have been associated with dysfunctional processing of facial expressions^28, 73, 74^, but also their impairment prevented individuals to integrate emotionally-salient facial features with gaze information about the target of the emotional response^75^. This was interpreted in line with appraisal models of emotion, whereby regions like amygdala and temporal pole elaborate emotional cues in relation to contextual information, and their relevance for the people involved^76^. Our data extend these previous findings, by revealing how these structures are necessary for adequate appraisal of others’ emotions from contextual information, without generalizing to similar evaluations outside the affective domain.

**Fig. 6:**
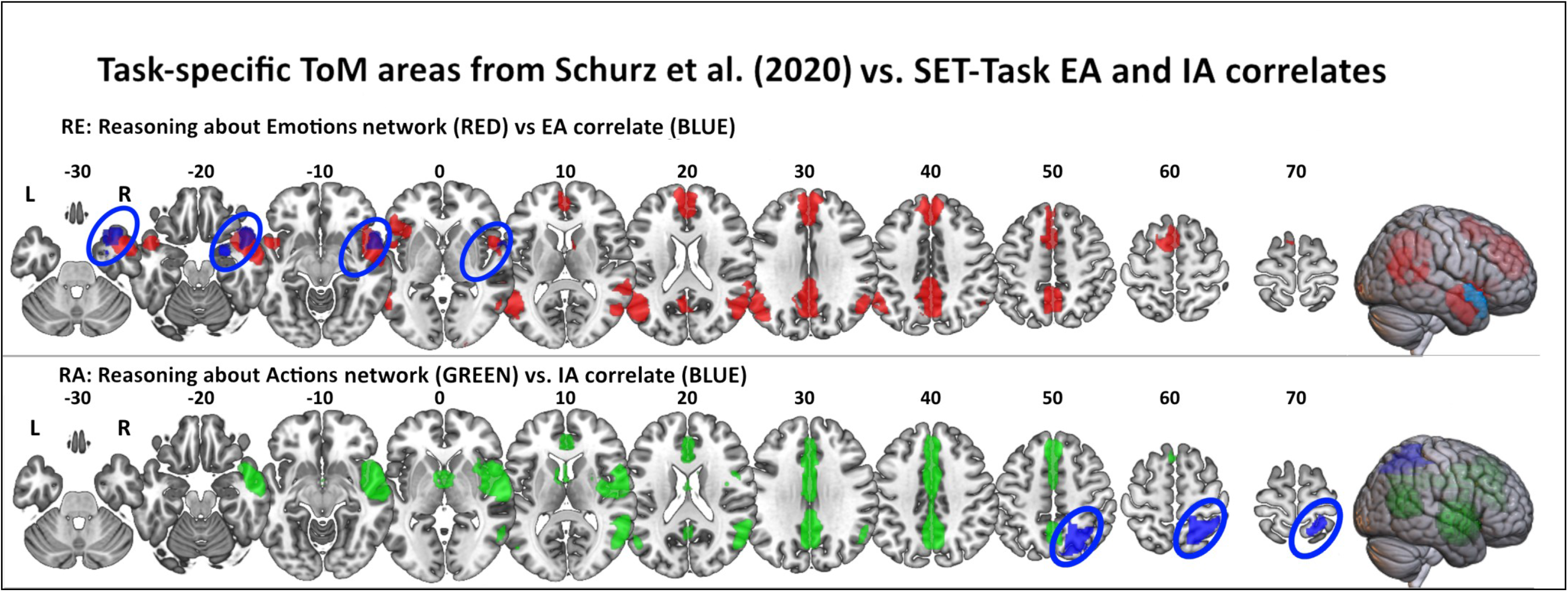
Superimposition of PLSM results to the ToM maps from Schurz et al. (2020) expected for the SET-Task conditions. Good overlap is found between SET-EA (Emotion Attribution) condition and the expected areas for Reasoning about Emotion (RE) ToM-type of task. PLSM results for SET-IA (Intention Attribution: right SPL) were however only adjacent but not overlapping with either TPJ or Precuneus, as expected for the Reasoning about Actions (RA) ToM-type of task.

#### Intention Attribution (Cognitive ToM)

Regarding intention attribution condition, this was selectively impaired following parietal lobe damage, and, more specifically, after damage to the right superior parietal cortex, when contrasting IA scores against EA (but not against the control CI condition). This effect is strongly reminiscent of that of Cohen-Zimerman et al.^48^ which mapped cognitive ToM deficits on the border between superior and inferior parietal cortex (albeit in the left hemisphere) and dorsolateral prefrontal cortex, after traumatic brain injuries. Our results converge with, but also extend, the findings from this previous research by revealing how lesion in the parietal cortex selectively influenced the ability to assess cognitive states and did not extend to the evaluation of emotions.

It could be argued that parietal portion implicated in cognitive ToM (both here and in Cohen-Zimerman et al.^48^) is part of the broad temporo-parietal region identified by functional neuroimaging studies testing cognitive social inferential abilities (see e.g.^8–11^ for meta-analyses). In this perspective, our data could be used as evidence of a convergence between neural correlates obtained across different approaches and populations. We suggest caution in this interpretation, for two reasons. First, lesion pattern from the present research is located in a clearly superior section of the parietal cortex than the TPJ activation map from neuroimaging research (see, Fig.6, for a superimposition of our data with the most recent meta-analysis of Schurz et al.,^10^). Second, neuroimaging studies in the typical brain clearly implicate TPJ, not only in cognitive ToM, but also in the appraisal of others’ emotions^14–16, 77^, at the point that the meta-analytic model from Schurz et al.^10^ describes this region also as part of a hybrid network where cognitive and affective inferential abilities co-exist. In this perspective, it has been argued that TPJ represents a key structure for a mentalistic strategy during emotion inference, according to which representations of people’s ongoing beliefs/thoughts/goals play a crucial role also in the inference of affective states (e.g., to understand if someone is sad, I need to have a clear idea about his/her thoughts^43^). This pattern clearly differs from our study, which documents the parietal cortex selectively implicated in IA (relative to EA). In this perspective, our data do not necessarily exclude that damage to TPJ might lead to ToM impairment, as already shown in single case analyses^24, 27^, one group study^28^ and researches employing neurostimulation techniques on typical brain^78, 79^. However, TPJ is not involved in the parietal cluster described in Figure 6, and its’ role in our study could have gone undetected, possibly due to limited number of patients with damage on this specific region. Indeed, it is worth noting that, differently from other regions of the brain left parietal lesions were less frequent (see Table 1 and Supplementary Fig.1). For example Cohen-Zimerman and colleagues^48^ identified a region at the border between superior and inferior parietal lobe (but on the *left*) as being associated to worse Cognitive ToM performance. A dataset with larger number of patients with parietal damage might allow to overcome this limitation.

To the best of our knowledge, the superior parietal involvement in IA could be interpreted in two possible ways. First, and most likely, the parietal cortex is indeed implicated in representing others’ cognitive states and, consequently, does also contribute in processing others’ emotion through a mentalistic strategy. However, damage to this region would not necessarily lead to an EA impairment as emotions could be inferred also through other (non-mentalistic) information, such as overt behaviour (e.g., trembling; see example in Figure 1), or contextual cues (the presence of a skull). Hence, neural structures located in the limbic system might compensate to the ToM impairment allowing to represent the others’ affect through an alternative strategy than accessing the characters’ thoughts/beliefs (see^14^ for a similar argument). Second, it is also possible that the parietal cortex processes a specific component of cognitive ToM that is not shared with EA.

Previous studies argued that *Cognitive* (but not *Affective*) ToM triggers regulatory processes for the inhibition of one’s own point of view, at the advantage of that of the story’s protagonist^25, 43^. Such mechanism has been previously associated with damage to the dlPFC^25, 43^, and is assumed to be particularly pronounced in those paradigms (like the False Beliefs task) where the distinction between the two viewpoints is made explicit. However, this interpretation might not apply to our case, as the SET paradigm from our study requires to infer a character’s intentions without any conflict with participants’ own point of view, and this might explain the lack of involvement of frontal lobe structures in our results. Alternatively, the effects observed in the parietal cortex could be specific for intentions (rather than cognitive states in general) and therefore reflect structures involved for action processing and understanding. Future research manipulating different kinds of cognitive states (intentions, beliefs, etc.) might shed more light on this issue.

### Limitations of the study and concluding remarks

In this study, we took great care to choose a clinically validated ToM task^53^, which was feasible for patients, and minimally biased by linguistic, memory, or attentional confounds. The drawback of this approach is that our data cannot be easily compared with those from previous research who employed more complex (lengthy and/or text-based) settings. Neuroimaging studies on typical brains have already showed a degree of variability in ToM network based on the paradigm employed^10^, and it is likely that the same applies also on lesion data. Furthermore, the brevity of the SET prevented us to investigate all aspects and facets of ToM, limiting ourselves to a gross distinction between cognitive, affective and non-social aspects of the task, without further exploring potential subdivisions within each category (intentions vs. beliefs; different kinds of contextual evaluations in emotional inference; etc.). Thus the present results (as those of many previous studies) might still be dependent on the task used. Future studies would be critical to replicate these findings also with other ToM tasks.

Furthermore, although the SET task is a clinically validated tool to detect ToM deficits with respect to the healthy population, what still needs to be assessed is *to what extent* ToM deficits are correlated to real-life everyday dysfunctional behaviours and on this aspect the neuropsychological literature on ToM is still largely insufficient^80^. Further research is thus needed on this regard.

Finally, this study benefited from a large sample of brain tumor patients, which allow for a widespread coverage of a large portion of the brain (see Supplementary Figure 1), including both lateral and medial structures, highly relevant for the investigation of ToM deficits. However, the heterogeneity of the tumour types and the effects of surgery might act as confounds, as suggested in recent review of ToM studies^80^. Brain reorganization is particularly pronounced in patients with LGG, and consequently potential lesion-symptom associations might go undetected, while Meningiomas are extra axial lesions producing milder and potentially more limited interference over brain functioning. However, the lack of an effect of tumor type over the results adds further confidence that the effects described are not linked to any specific type of interaction between the different tumors (and their biological behavior) and the brain tissue, but rather the specific location in which the lesion occurs. Hence, although we cannot exclude that factors like reorganization/pressure might be at play in our sample, we are reasonably confident that their impact on the ToM scores should be negligible.

Notwithstanding these limitations, this study provides new evidence on the functional properties of the network mediating ToM deficits. Part of this network (frontal cortex) contributes to ToM abilities in a non-selective fashion, through the same inferential mechanism involved in making causal judgments, and modulated by executive function abilities. Other portions of the network are selectively involved in ToM, with the parietal cortex underlying deficits in inferring others’ intentions, and temporal pole and amygdala implicated instead in emotion attribution. This pattern is only in partial agreement with the findings from neuroimaging literature, but overall, our data provide new evidence of segregation in brain network necessary for human ToM abilities.

## SUPPLEMENTARY MATERIALS

### Detailed SET Task description

In SET task patients are required to select the correct ending of a comic strip composed of three vignettes, depicted in the upper row of a sheet. In the lower row of the same sheet, three further vignettes (possible alternative endings) are provided (see Fig.1). The task consists of two Theory of Mind conditions, i.e., Intention Attribution (IA) and Emotion Attribution (EA), in which the inference of the correct ending of the story is based upon the correct interpretation of the intention or the emotional state of the main character respectively. In a further control condition (Causal Infererence, CI) the correct answer is based on the inference of causal relations between events, based on the knowledge of physical properties of the world, thus allowing to control for more general impairments in basic reasoning skills. The number of trials per each condition is of 6. Thus, overall there are 18 trials in the task. Answers are scored as correct (1) or incorrect (0) according to whether the correct alternative is chosen or not. There is no time limit for responding; if the subject, however, was unable to answer, he/she was encouraged to guess. No feedback is provided but, prior to the task, participants underwent run-in practice trial in which the task is operatively explained and the correct answer revealed in case of error. The participant is invited to assure he/she understood the meaning of the comic strip and every possible doubt or misunderstanding is clarified before the beginning of the true task session. The subject was moreover encouraged to describe the scene aloud and imagine the conclusion every time he/she experienced difficulties.

### Sensitivity analysis for the ANCOVA

As explained in the main text, we run an ANCOVA to unveil effects of the within-subject factor “ToM Condition”, either alone or in interaction with the lesion location (grouping factor “Lobe Location” and “Hemisphere”) while accounting for the influence of nuisance grouping variables. As a preliminary step, we run a sensitivity analysis assessing the minimum effect size observable by our sample with the present model, at power = 0.80 and α = 0.05. The analysis was carried out using G*Power 3.1.9.7 software, under the “generic F-test” which allows the highest flexibility in the design structure, by estimating a non-centrality parameter λ only based on the degrees of freedom of the effect of interest. The resulting λ was then converted into an estimate of the effect sizes based on the following formula^#^ 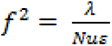 where

*N =* sample size

*f =* effect size (Cohen’s *f*)

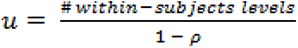

*ρ =* assumed correlation between the within-subject measures (set at 0.5, consistently with G*Power default parameters).

*ε =* non-sphericity correction parameter (here set to 1)

The resulting Cohen’s f was in turn, can be reformulated in terms of eta-square: 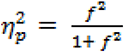.

This analysis revealed that our ANCOVA could reliably detect effects associated with the “ToM Condition” within-subjects factor of at least 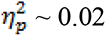.

### Sensitivity analysis at the anatomical level (PLSM analysis)

We run a sensitivity analysis also for the parcel-based lesion mapping (PLSM). In this model, the brain is divided into 85 regions of interest (ROIs) based on the AAL atlas (regions damaged in less than 6 patients were excluded; see methods), and the lesion-behavior relationship is estimated through linear models where the task score is tested as function of the percentage in which each ROI is damaged [from 0 to 100%]. As first step, therefore run sensitivity analysis for correlation coefficients (as available on G*Power software) to establish the minimum effect size detectable by the present sample (N = 105) with power = 0.80 and α = 0.05/85 (corresponding to a Bonferroni-corrected α = 0.05, with multiple comparisons involving all 85 ROIs). We found that the present design was powerful enough to detect effects of at least r = 0.38 in any ROI. Very similar results (r = 0.37) are obtained under a less conservative α = 0.001 (uncorrected).

The latter sensitivity analysis relies however exclusively on sample size, without taking into account lesion distribution. It could be argued that, in some ROI, sensitivity might be affected by the lack of patients with lesion in that area, which leads to predictors characterized by many 0s, and only a few with some damage. To tackle this problem, we adapted the sensitivity analysis to takes into account the idiosyncratic damage distribution of each ROI. Therefore a complementary Monte Carlo simulation was run in which, for each ROI, 5000 behavioral scores were simulated that correlated with the actual lesion data with a known effect size. Each of these simulations was assessed statistically through rigorous permutation analyses (5000 data shufflings) for unbiased estimation of the null distribution (as it was implemented in the main analysis). Through an iterative approach, the minimum effect size that could be observed with power 0.8 and α = 0.05/85 was identified. Across the 85 regions, the minimum effect size ranged between *r* = [0.37-0.39], with marginal fluctuations between the different ROIs and strong agreement with the analysis carried out with G*Power. Very similar effects (*r* = [0.37-0.38]) were observable under a less conservative α = 0.001 (uncorrected). Please find enclosed the ad hoc script in which the simulation was run.

---

**Supplementary Figure 1:**
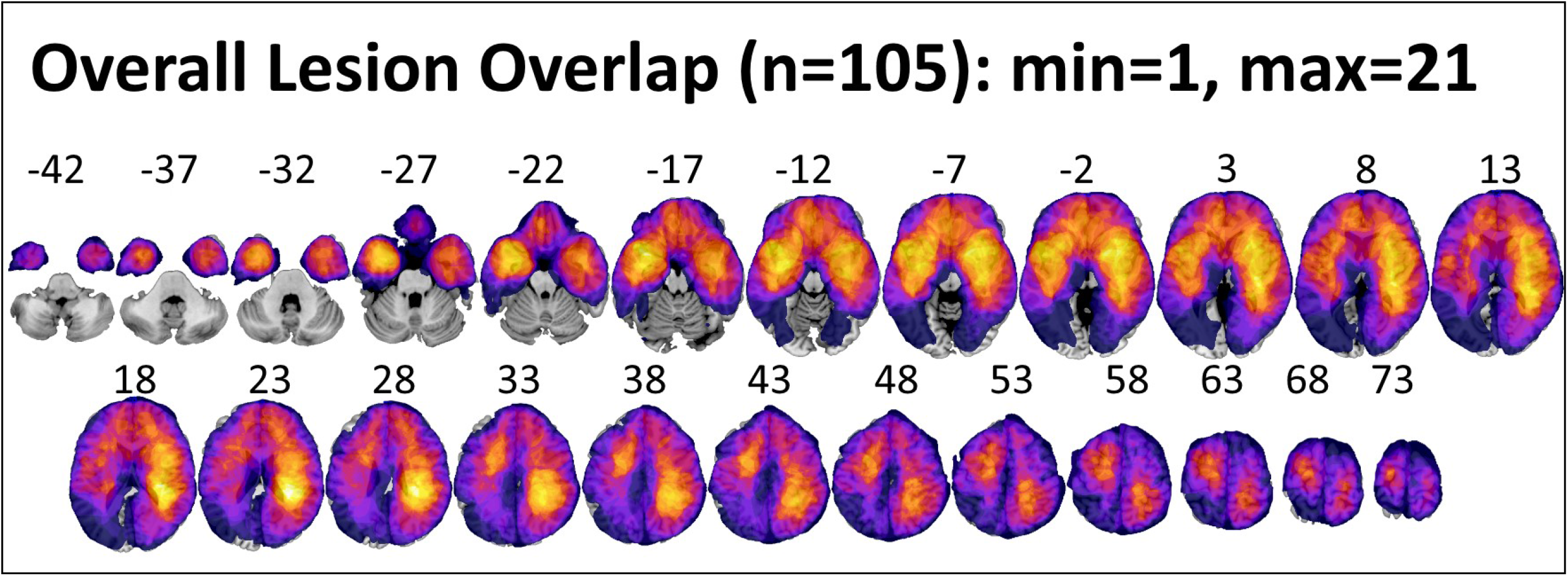
Lesion overlap maps showing anatomical coverage of lesions included in the behavioral analysis.

**Supplementary Table 1:**
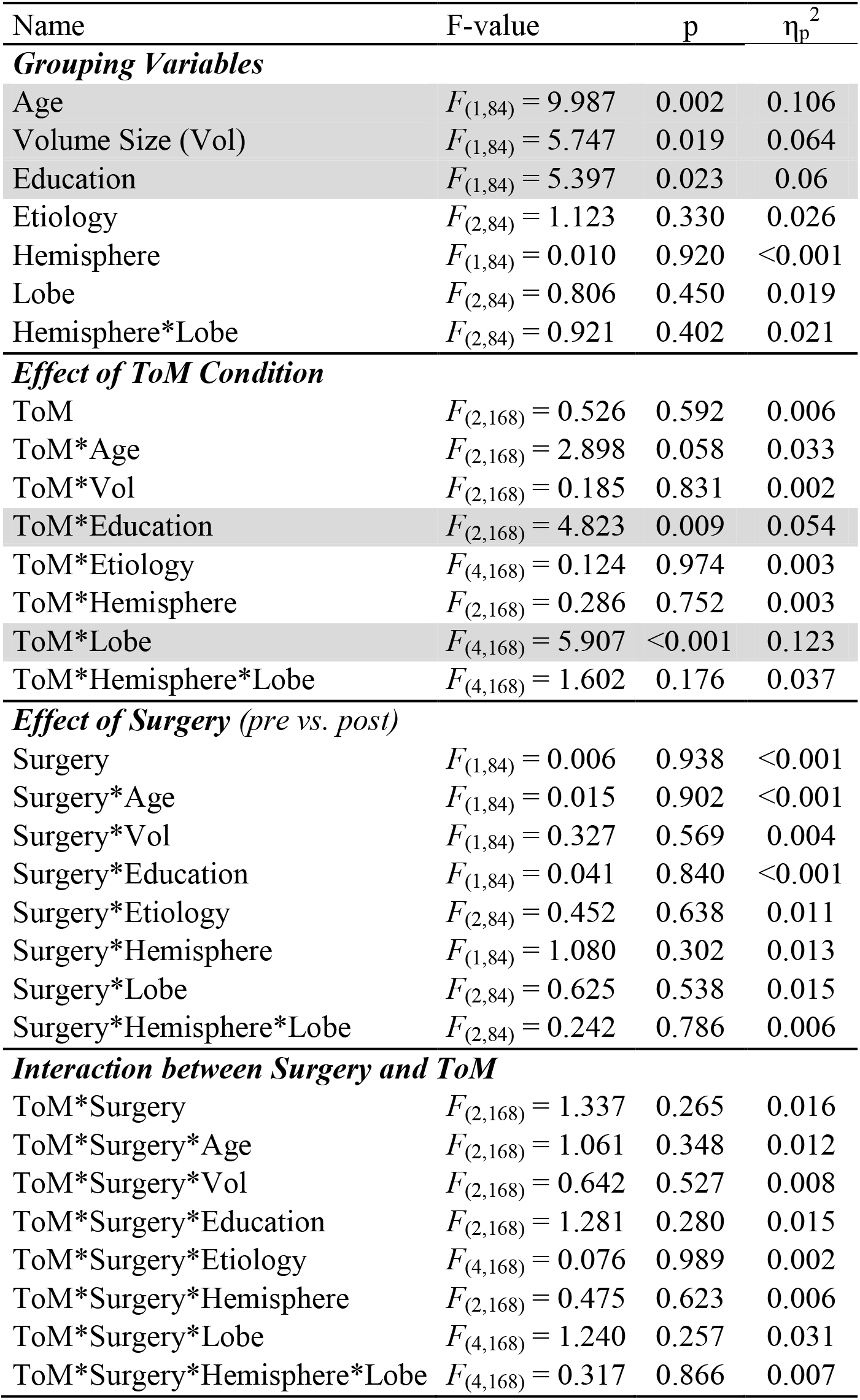
complete report of ANCOVA results controlling for the potential effects of age, education and tumor type over the three ToM conditions of the SET Task. In grey significant effects. A highly significant ToM condition x Location interaction was found despite the influence of the covariates variables considered. Tumor type (Etiology) did not have any effect at any level. No effects were also found for surgery at any level.

**Supplementary Table 2:**
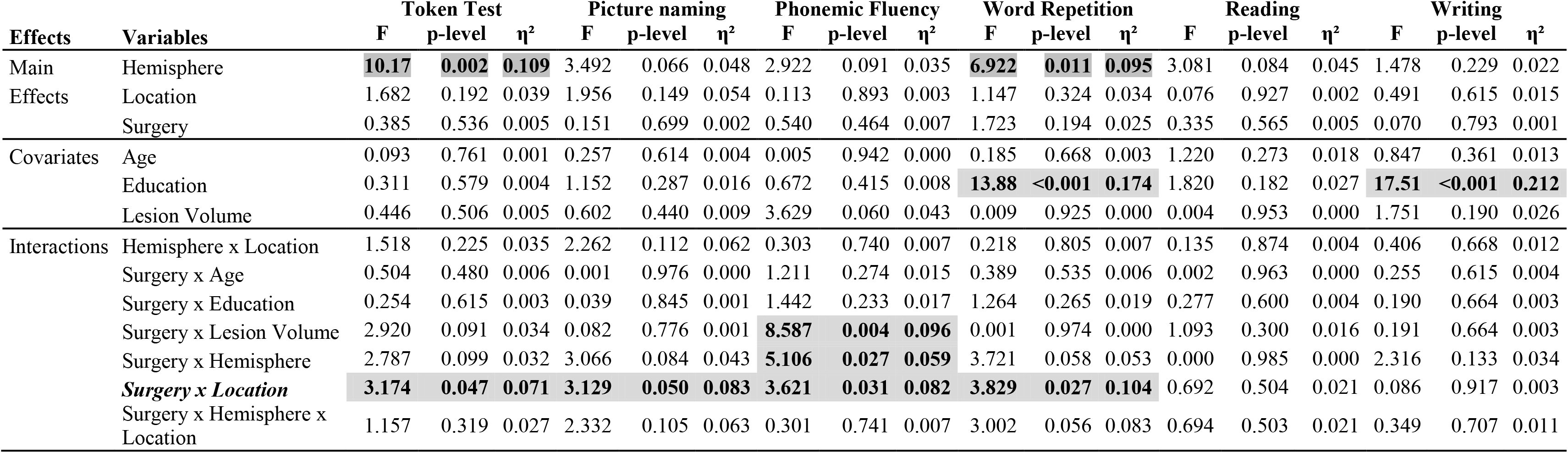
Assessment of surgery effects in baseline language tasks. Baseline language tasks used for the routine pre-post surgery assessment of language skills of patients, do not have an alternative version to be used after the short test-retest delay (about a week), similarly to the SET-Task used to investigate ToM skills. However, even when repeating the same task after only a week, no learning effect is evident in the performance of patients and clear detrimental immediate surgery effects are present as clearly shown by the significant Surgery x Location interaction found for 4 out of 6 of the language tasks considered. Post-hoc analysis (Tukey Test) of the significant interactions shows that in 3/4 cases Temporal lobe patients alone (all p<.004) had worse language performance immediately after surgery (for Phonemic Fluency p=.080).

**Supplementary Table 3:**
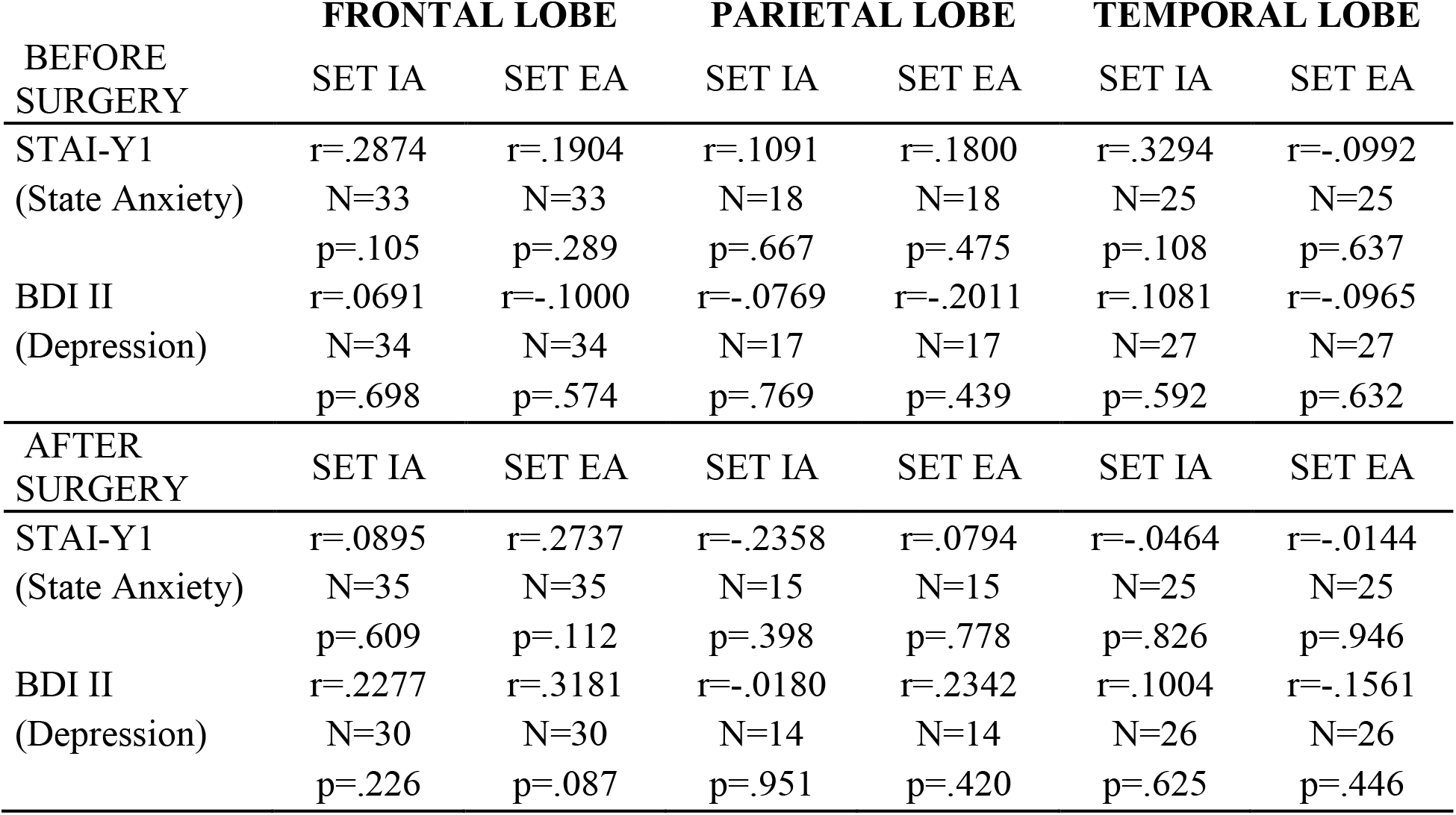
Correlation matrix to assess potential correlations between basic mood state (State Anxiety – STAI-Y1 z-scores; Depression – BDI-II percentile scores) and Theory of Mind z-scores. No significant relation was found either before or after surgery. P-levels reported are Bonferroni uncorrected.

## Notes

### Competing Interest Statement

The authors have declared no competing interest.

