## Supplementary material for "Cognitive and affective Theory of Mind double dissociation after parietal and temporal lobe tumors": Sensitivity_Analysis_PLSM.m

```
function myfunction
%
% Function for the assessment of the minimum sample size in a PLSM (as implemented in NiiStat)
%
% parameters
%
nsamples = 5000; % Number of Monte Carlo simulations and permutation reshuffles (same value for both) 
des_pow = 0.80;  % desired power
alpha = 0.001;   % desired alpha level
%
% the script is designed to work on the "power.mat" file provided by NiiStat software after each analysis.
% This file contains the relevant variable "les"
% "les" = Subject x ROI matrix, containing the relative amount of damage of each patient for each ROI on a scale from 0 (undamaged ROI) to 1 (entrirely damaged ROI). 
%
m = load('power.mat');
nVox = size(m.les,2);
pow = NaN(1,nVox);
%
% loop for voxel
%
parfor ivox = 1: nVox
    pow(ivox) = ccd_roivox(ivox,alpha,nsamples,des_pow,m);
end
%
% output file: 1 x ROI vector, containing the minimum effect size (in r) for each ROI
%
save power_result.mat pow -V6

function temppow = ccd_roivox(ivox,alpha,nsamples,des_pow,m)
N = size(m.les,1);
tempincr = 0.1;
tpow2 = 1;
rcoeff = 0.2;
rcoeff2 = NaN;
%
% start the iterative process
%
while tpow2 > des_pow + 0.005
    tpow = 0;
    while tpow < des_pow
        rcoeff = rcoeff + tempincr;
        %
        % simulate correlations at a given r
        %
        L = chol([1 rcoeff; rcoeff 1]);
        r = zeros(1,nsamples);
        pv = zeros(1,nsamples);
        for j = 1:nsamples
            xy = normrnd(mean(m.les(:,ivox)),std(m.les(:,ivox)),N,1);
            xy(:,2) = m.les(:,ivox);
            xy = xy*L;
            opt = permcorr(xy(:,1),xy(:,2),0.05,nsamples,1,'Pearson');
            r(j) = opt.r;
            pv(j) = opt.pval;
        end
        tpow = binofit(sum(pv<alpha),nsamples);
    end
    rcoeff2 = rcoeff;
    rcoeff = rcoeff - tempincr;
    tempincr = tempincr/3;
    tpow2 = tpow;
end
%
temppow = rcoeff2;
disp(['ROI n. ',num2str(ivox),'; # lesioned patients ',num2str(sum(m.les(:,ivox)>0)),'; power = ',num2str(tpow),'; min effect size, r = ',num2str(rcoeff2)])

function outp = permcorr(x,y,pval,npermut,tail,rtype)
%
% subfunction for permutation analysis of correlations
%
% outp = permcorr(x,y,pval,npermut,tail)
%
% x & y = 2 vectors
% pval = pvalue (usually 0.05)
% npermut = Number of permutations (usually 5000)
% tail = 2 ---> bidirectional; 1 ----> higher; -1 -----> lower 
% rtype = 'Pearson','Spearman','Kendall'
%

%
% seed the random number generator
%
rand('state',sum(100*clock));
%
% calculate REAL effect
%
if size(x,1) == 1
	x = x';
end
if size(y,1) == 1
	y = y';
end
r = corr([x,y],'rows','pairwise','type',rtype);
outp.r = r(1,2);
%
% create shuffled data
%
clear remr x_max y_max r
x_mat = repmat(x,1,npermut);
y_mat = repmat(y,1,npermut);
for iperm = 1:npermut
    x_mat(:,iperm) = x_mat(randperm(length(x)),iperm);
    y_mat(:,iperm) = y_mat(randperm(length(x)),iperm);
end
r = corr(x_mat,y_mat,'rows','pairwise','type',rtype);
remr = diag(r);
%
% calculate ctoff
%
if tail == 2
    outp.ctoff(1) = prctile(remr,pval/2*100);
    outp.ctoff(2) = prctile(remr,(1-pval/2)*100);
    outp.pval = (sum(remr >= abs(outp.r)) + sum(remr <= -abs(outp.r)))/npermut;
    if outp.r < outp.ctoff(1)
        outp.h='*';
    elseif outp.r > outp.ctoff(2)
        outp.h='*';
    else
        outp.h=' ';
    end
    disp(['real effect: ', num2str(outp.r),' [', num2str(outp.ctoff(1)),', ', num2str(outp.ctoff(2)),'], p = ',num2str(outp.pval),outp.h]);
elseif tail == 1
    outp.ctoff = prctile(remr,(1-pval)*100);
    outp.pval = (sum(remr >= outp.r))/npermut;
    if outp.r > outp.ctoff
        outp.h='*';
    else
        outp.h=' ';
    end
    %disp(['real effect: ', num2str(outp.r),' [', num2str(outp.ctoff),'], p = ',num2str(outp.pval),outp.h]);
elseif tail == -1
    outp.ctoff = prctile(remr,(pval)*100);
    outp.pval = (sum(remr <= outp.r))/npermut;
    if outp.r < outp.ctoff
        outp.h='*';
    else
        outp.h=' ';
    end
    disp(['real effect: ', num2str(outp.r),' [', num2str(outp.ctoff),'], p = ',num2str(outp.pval),outp.h]);
end
```
