## Supplementary material for "Cognitive and affective Theory of Mind double dissociation after parietal and temporal lobe tumors": SUPPLEMENTARY MATERIALS FOR PREPRINT.pdf

*N* = sample size
*f* = effect size (Cohen's *f*)
$u = \frac{\#\ within\text{−}subjects\ levels}{1 - \rho}$
$\rho$ = assumed correlation between the within-subject measures (set at 0.5, consistently with G*Power default parameters).
$\varepsilon$ = non-sphericity correction parameter (here set to 1)

The resulting Cohen's f was in turn, can be reformulated in terms of eta-square: $\eta_p^2 = \frac{f^2}{1+ f^2}$.

This analysis revealed that our ANCOVA could reliably detect effects associated with the "ToM Condition" within-subjects factor of at least $\eta_p^2 \sim 0.02$.

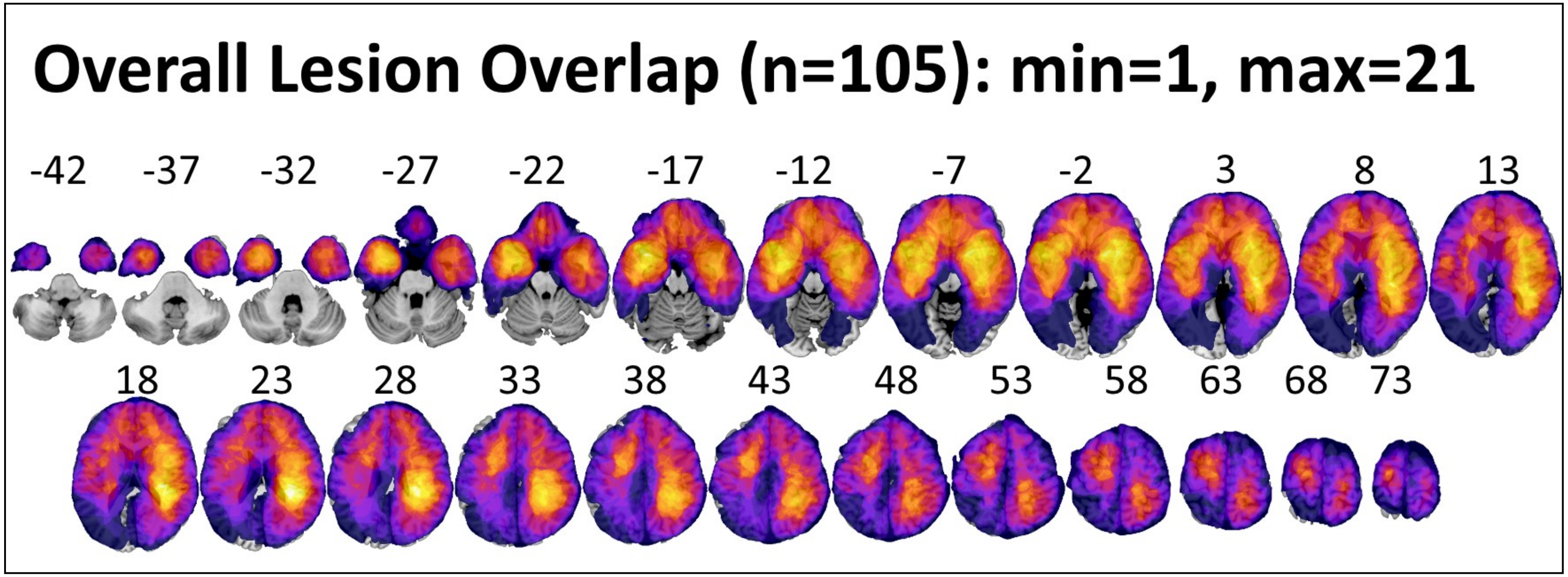


Supplementary Figure 1: Lesion overlap maps showing anatomical coverage of lesions included in the behavioral analysis.

| Name | F-value | p | $\eta_p^2$ |
|---|---|---|---|
| ***Grouping Variables*** | | | |
| Age | $F_{(1,84)} = 9.987$ | 0.002 | 0.106 |
| Volume Size (Vol) | $F_{(1,84)} = 5.747$ | 0.019 | 0.064 |
| Education | $F_{(1,84)} = 5.397$ | 0.023 | 0.06 |
| Etiology | $F_{(2,84)} = 1.123$ | 0.330 | 0.026 |
| Hemisphere | $F_{(1,84)} = 0.010$ | 0.920 | <0.001 |
| Lobe | $F_{(2,84)} = 0.806$ | 0.450 | 0.019 |
| Hemisphere*Lobe | $F_{(2,84)} = 0.921$ | 0.402 | 0.021 |
| ***Effect of ToM Condition*** | | | |
| ToM | $F_{(2,168)} = 0.526$ | 0.592 | 0.006 |
| ToM*Age | $F_{(2,168)} = 2.898$ | 0.058 | 0.033 |
| ToM*Vol | $F_{(2,168)} = 0.185$ | 0.831 | 0.002 |
| ToM*Education | $F_{(2,168)} = 4.823$ | 0.009 | 0.054 |
| ToM*Etiology | $F_{(4,168)} = 0.124$ | 0.974 | 0.003 |
| ToM*Hemisphere | $F_{(2,168)} = 0.286$ | 0.752 | 0.003 |
| ToM*Lobe | $F_{(4,168)} = 5.907$ | <0.001 | 0.123 |
| ToM*Hemisphere*Lobe | $F_{(4,168)} = 1.602$ | 0.176 | 0.037 |
| ***Effect of Surgery*** *(pre vs. post)* | | | |
| Surgery | $F_{(1,84)} = 0.006$ | 0.938 | <0.001 |
| Surgery*Age | $F_{(1,84)} = 0.015$ | 0.902 | <0.001 |
| Surgery*Vol | $F_{(1,84)} = 0.327$ | 0.569 | 0.004 |
| Surgery*Education | $F_{(1,84)} = 0.041$ | 0.840 | <0.001 |
| Surgery*Etiology | $F_{(2,84)} = 0.452$ | 0.638 | 0.011 |
| Surgery*Hemisphere | $F_{(1,84)} = 1.080$ | 0.302 | 0.013 |
| Surgery*Lobe | $F_{(2,84)} = 0.625$ | 0.538 | 0.015 |
| Surgery*Hemisphere*Lobe | $F_{(2,84)} = 0.242$ | 0.786 | 0.006 |
| ***Interaction between Surgery and ToM*** | | | |
| ToM*Surgery | $F_{(2,168)} = 1.337$ | 0.265 | 0.016 |
| ToM*Surgery*Age | $F_{(2,168)} = 1.061$ | 0.348 | 0.012 |
| ToM*Surgery*Vol | $F_{(2,168)} = 0.642$ | 0.527 | 0.008 |
| ToM*Surgery*Education | $F_{(2,168)} = 1.281$ | 0.280 | 0.015 |
| ToM*Surgery*Etiology | $F_{(4,168)} = 0.076$ | 0.989 | 0.002 |
| ToM*Surgery*Hemisphere | $F_{(2,168)} = 0.475$ | 0.623 | 0.006 |
| ToM*Surgery*Lobe | $F_{(4,168)} = 1.240$ | 0.257 | 0.031 |
| ToM*Surgery*Hemisphere*Lobe | $F_{(4,168)} = 0.317$ | 0.866 | 0.007 |

Supplementary table 1: complete report of ANCOVA results controlling for the potential effects of age, education and tumor type over the three ToM conditions of the SET Task. In grey significant effects. A highly significant ToM condition x Location interaction was found despite the influence of the covariates variables considered. Tumor type (Etiology) did not have any effect at any level. No effects were also found for surgery at any level.

| Effects | Variables | Token Test | | | Picture naming | | | Phonemic Fluency | | | Word Repetition | | | Reading | | | Writing | | |
|---|---|---|---|---|---|---|---|---|---|---|---|---|---|---|---|---|---|---|---|
| | | F | p-level | η² | F | p-level | η² | F | p-level | η² | F | p-level | η² | F | p-level | η² | F | p-level | η² |
| Main Effects | Hemisphere | ***10.17*** | ***0.002*** | ***0.109*** | 3.492 | 0.066 | 0.048 | 2.922 | 0.091 | 0.035 | 6.922 | 0.011 | 0.095 | 3.081 | 0.084 | 0.045 | 1.478 | 0.229 | 0.022 |
| | Location | 1.682 | 0.192 | 0.039 | 1.956 | 0.149 | 0.054 | 0.113 | 0.893 | 0.003 | 1.147 | 0.324 | 0.034 | 0.076 | 0.927 | 0.002 | 0.491 | 0.615 | 0.015 |
| | Surgery | 0.385 | 0.536 | 0.005 | 0.151 | 0.699 | 0.002 | 0.540 | 0.464 | 0.007 | 1.723 | 0.194 | 0.025 | 0.335 | 0.565 | 0.005 | 0.070 | 0.793 | 0.001 |
| Covariates | Age | 0.093 | 0.761 | 0.001 | 0.257 | 0.614 | 0.004 | 0.005 | 0.942 | 0.000 | 0.185 | 0.668 | 0.003 | 1.220 | 0.273 | 0.018 | 0.847 | 0.361 | 0.013 |
| | Education | 0.311 | 0.579 | 0.004 | 1.152 | 0.287 | 0.016 | 0.672 | 0.415 | 0.008 | **13.88** | **<0.001** | **0.174** | 1.820 | 0.182 | 0.027 | **17.51** | **<0.001** | **0.212** |
| | Lesion Volume | 0.446 | 0.506 | 0.005 | 0.602 | 0.440 | 0.009 | 3.629 | 0.060 | 0.043 | 0.009 | 0.925 | 0.000 | 0.004 | 0.953 | 0.000 | 1.751 | 0.190 | 0.026 |
| Interactions | Hemisphere x Location | 1.518 | 0.225 | 0.035 | 2.262 | 0.112 | 0.062 | 0.303 | 0.740 | 0.007 | 0.218 | 0.805 | 0.007 | 0.135 | 0.874 | 0.004 | 0.406 | 0.668 | 0.012 |
| | Surgery x Age | 0.504 | 0.480 | 0.006 | 0.001 | 0.976 | 0.000 | 1.211 | 0.274 | 0.015 | 0.389 | 0.535 | 0.006 | 0.002 | 0.963 | 0.000 | 0.255 | 0.615 | 0.004 |
| | Surgery x Education | 0.254 | 0.615 | 0.003 | 0.039 | 0.845 | 0.001 | 1.442 | 0.233 | 0.017 | 1.264 | 0.265 | 0.019 | 0.277 | 0.600 | 0.004 | 0.190 | 0.664 | 0.003 |
| | Surgery x Lesion Volume | 2.920 | 0.091 | 0.034 | 0.082 | 0.776 | 0.001 | **8.587** | **0.004** | **0.096** | 0.001 | 0.974 | 0.000 | 1.093 | 0.300 | 0.016 | 0.191 | 0.664 | 0.003 |
| | Surgery x Hemisphere | 2.787 | 0.099 | 0.032 | 3.066 | 0.084 | 0.043 | **5.106** | **0.027** | **0.059** | 3.721 | 0.058 | 0.053 | 0.000 | 0.985 | 0.000 | 2.316 | 0.133 | 0.034 |
| | ***Surgery x Location*** | **3.174** | **0.047** | **0.071** | **3.129** | **0.050** | **0.083** | **3.621** | **0.031** | **0.082** | **3.829** | **0.027** | **0.104** | 0.692 | 0.504 | 0.021 | 0.086 | 0.917 | 0.003 |
| | Surgery x Hemisphere x Location | 1.157 | 0.319 | 0.027 | 2.332 | 0.105 | 0.063 | 0.301 | 0.741 | 0.007 | 3.002 | 0.056 | 0.083 | 0.694 | 0.503 | 0.021 | 0.349 | 0.707 | 0.011 |

Supplementary Table 2: Assessment of surgery effects in baseline language tasks. Baseline language tasks used for the routine pre-post surgery assessment of language skills of patients, do not have an alternative version to be used after the short test-retest delay (about a week), similarly to the SET-Task used to investigate ToM skills. However, even when repeating the same task after only a week, no learning effect is evident in the performance of patients and clear detrimental immediate surgery effects are present as clearly shown by the significant Surgery x Location interaction found for 4 out of 6 of the language tasks considered. Post-hoc analysis (Tukey Test) of the significant interactions shows that in 3/4 cases Temporal lobe patients alone (all $p<.004$) had worse language performance immediately after surgery (for Phonemic Fluency $p=.080$).

| BEFORE SURGERY | FRONTAL LOBE | | PARIETAL LOBE | | TEMPORAL LOBE | |
|---|---|---|---|---|---|---|
| | SET IA | SET EA | SET IA | SET EA | SET IA | SET EA |
| STAI-Y1 | r=.2874 | r=.1904 | r=.1091 | r=.1800 | r=.3294 | r=-.0992 |
| (State Anxiety) | N=33 | N=33 | N=18 | N=18 | N=25 | N=25 |
| | p=.105 | p=.289 | p=.667 | p=.475 | p=.108 | p=.637 |
| BDI II | r=.0691 | r=-.1000 | r=-.0769 | r=-.2011 | r=.1081 | r=-.0965 |
| (Depression) | N=34 | N=34 | N=17 | N=17 | N=27 | N=27 |
| | p=.698 | p=.574 | p=.769 | p=.439 | p=.592 | p=.632 |
| AFTER SURGERY | SET IA | SET EA | SET IA | SET EA | SET IA | SET EA |
| STAI-Y1 | r=.0895 | r=.2737 | r=-.2358 | r=.0794 | r=-.0464 | r=-.0144 |
| (State Anxiety) | N=35 | N=35 | N=15 | N=15 | N=25 | N=25 |
| | p=.609 | p=.112 | p=.398 | p=.778 | p=.826 | p=.946 |
| BDI II | r=.2277 | r=.3181 | r=-.0180 | r=.2342 | r=.1004 | r=-.1561 |
| (Depression) | N=30 | N=30 | N=14 | N=14 | N=26 | N=26 |
| | p=.226 | p=.087 | p=.951 | p=.420 | p=.625 | p=.446 |

Supplementary table 3: Correlation matrix to assess potential correlations between basic mood state (State Anxiety – STAI-Y1 z-scores; Depression – BDI-II percentile scores) and Theory of Mind z-scores. No significant relation was found either before or after surgery. P-levels reported are Bonferroni uncorrected.
